# A Bayesian method using sparse data to estimate penetrance of disease-associated genetic variants

**DOI:** 10.1101/571158

**Authors:** Brett M. Kroncke, Derek K. Smith, Andrew M. Glazer, Dan M. Roden, Jeffrey D. Blume

## Abstract

**Purpose:** A major challenge in genomic medicine is how to best predict risk of disease from rare variants discovered in Mendelian disease genes but with limited phenotypic data. We have recently used Bayesian methods to show that *in vitro* functional measurements and computational pathogenicity classification of variants in the cardiac gene *SCN5A* correlate with rare arrhythmia penetrance. We hypothesized that similar predictors could be used to impute variant-specific penetrance prior probabilities.

**Methods:** From a review of 756 publications, we developed a pattern mixture algorithm, based on a Bayesian Beta-Binomial model, to generate *SCN5A* variant-specific penetrance priors for the heart arrhythmia Brugada syndrome (BrS).

**Results:** The resulting priors correlate with mean BrS penetrance posteriors (cross validated R^2^= 0.41). *SCN5A* variant function and structural context provide the most information predictive of BrS penetrance. The resulting priors are interpretable as equivalent to the observation of affected and unaffected carriers.

**Conclusions:** Bayesian estimates of penetrance can efficiently integrate variant-specific data (e.g. functional, structural, and sequence) to accurately estimate disease risk attributable to individual variants. We suggest this formulation of penetrance is quantitative, probabilistic, and more precise than, but consistent with, discrete pathogenicity classification approaches.

## Introduction

A major challenge to integrating genotype information into clinical care is accurately linking genetic variants to disease. As cheap whole genome, exome, and gene panel sequencing become more widely used, the genetics community is frequently observing novel, ultra-rare variants—ones carried by a single or few (often related) individuals. Indeed, *most* variants found in large genome sequencing efforts are novel or ultra rare;^1,2^ and the majority of these variants will never be observed in a sufficient number of carriers to ascertain a strong statistical association with disease. In addition, with recent large-scale genetic sequencing efforts taking place around the world, the genetics community is identifying greater numbers of individuals, ostensibly unaffected, who carry variants previously thought to be disease-inducing.^3,4^ As a consequence, both insufficient carrier counts and increasing conflict of annotations are causing many diagnostic laboratories that previously confidently annotated genetic variants as “Likely Pathogenic” or “Pathogenic”, to annotate increasingly more conservatively, now calling most variants “Variants of Uncertain Significance” (VUS).^5-8^

To help assess the significance of variants, the American College of Medical Genetics and Genomics (ACMG) suggests criteria to integrate population, functional, computational, and segregation data using several described heuristics to classify variants.^9,10^ While this classification yields a common scale by which variants can be interpreted and compared, for the majority of variants, only a subset of those data is known (computed or experimentally determined). Additionally, the annotation framework suggests classifications ranging from “pathogenic/likely pathogenic” to “likely benign/benign”, with variants not confidently placed into either of these categories classified as VUS, emphasizing the overall degree of uncertainty in classification instead of degree of pathogenicity. We suggest incorporating both degree of uncertainty and degree of pathogenicity will enable the increased use of genetics in medicine.

Bayesian methods are a promising approach to address the rare, novel variant annotation opportunity described above given their ability to estimate the likelihood of an outcome, even when data are sparse, by integrating multiple lines of evidence. In this study, we built a Bayesian model to estimate the probability a variant will cause a disease outcome; here, this probability is interpreted as the penetrance of the variant. Our goals of the analyses are: 1) to produce a high-quality estimate of penetrance that can be applied to variants for which carrier data are limited or unavailable; and 2) to develop a method for generating a data-driven prior distribution of penetrance which can be updated as new data become available for previously observed or unobserved variants. The first goal relies on statistical methods of “borrowing strength” or sharing information across variants. The second goal will produce variant-specific, quantitative penetrance priors—especially informative for rare variants—even in the absence of a large number of carriers.

We develop this framework for the rare cardiac arrhythmia disorder Brugada Syndrome (BrS), which is linked to rare loss-of-function variants in the cardiac sodium channel *SCN5A*.^11^ Our proposed approach can efficiently integrate functional and structural data, previously published variant classifiers, and carrier counts observed in affected and unaffected populations to estimate the BrS penetrance attributable to individual *SCN5A* variants. Additionally, our framework has the flexibility to incorporate more information (e.g. additional variant functional characteristics, carrier demographics, etc.) as more variant or carrier data are discovered. As part of this analysis, we produce a structure-based metric to quantitate enrichment in “hotspot” regions of the protein structure for variants with higher penetrance.^12^ We show herein that we can leverage the quantitative relationship between predictive features, determinable for all variants, to impute variant-specific priors for BrS penetrance. We suggest these methodologies can be extended to other genes and disorders in order to enable quantitative interpretation of variants, from rare to common, probabilistically and quantitatively.^13^

## Materials and Methods

These analyses focus on the *SCN5A* gene, where individual variants are known to influence the clinical presentation of the autosomal dominant arrhythmia Brugada Syndrome (BrS).^14^, ^15^ We define cases as individuals with either a spontaneous or drug-induced ECG BrS pattern.^16^ Penetrance is generally defined as the fraction of variant carriers who are BrS cases. We define an estimated BrS penetrance as the average variant-specific posterior penetrance, denoted as the following:

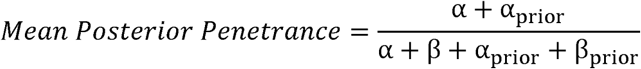

Where α is the number of BrS cases who are variant carriers and β is the number of unaffected carriers of a specific variant. As the total number of carriers increases, the estimated penetrance converges to the traditional definition. The mean posterior penetrance can be thought of as a shrunken estimate of the observed penetrance,^17^ especially for variants with small numbers of known carriers. The priors can have a large influence on the estimates, and how to select them is the focus of this paper.

To generate priors from our available data, we use a variation of the expectation maximization (EM) algorithm.^18^ Our modified EM algorithm is an iterative technique composed of two steps: 1) calculate the expected penetrance from an empirical Bayes penetrance model and 2) fit a regression model of our estimated penetrance on variant-specific characteristics by maximum likelihood:

Penetrance Estimate_i_

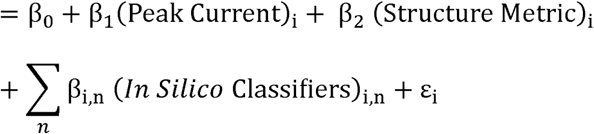

Here peak current is an *in vitro* measurement of the maximum current through a channel (normalized to wild type), structure metric is the penetrance density^12^ (detailed in the supplement), and *in silico* classifiers is a vector populated with commonly used variant classification servers such as PROVEAN and PolyPhen (see below); all predictors used are continuous, not categorical or binary. The fitted model is then used to generate an updated prior distribution and subsequent posterior expected penetrance and this process is iterated until it converges to the maximum likelihood solution (Figure 1). Resulting models can be used to generate a variant-specific predicted penetrance and nonparametric variance estimate based on local averaging. Using a beta-binomial model to estimate penetrance, the prior parameters (α_prior_ and β_prior_) are identifiable from a predicted penetrance and its associated variance. Note that we do not observe the actual penetrance for any given variant; the penetrance is a latent variable. For comparison, we generated predicted penetrance values using a standard empirical Bayes method which generated a single empirical prior for all variants (called empirical prior throughout the text, Figure S1). As a result, we compare our EM penetrance predictions to the posterior mean penetrance derived from the two separate priors: 1) prior derived by the weighted average penetrance over all variants in the dataset (empirical prior, not variant specific and a more conservative estimate) and 2) prior imputed by the EM algorithm using pattern mixture models to accommodate for variants with missing features. Additionally, we generated priors applying the full dataset or by removing one variant at a time from all stages of prior generation (leave one out cross validation; LOOCV).

**Figure 1.**
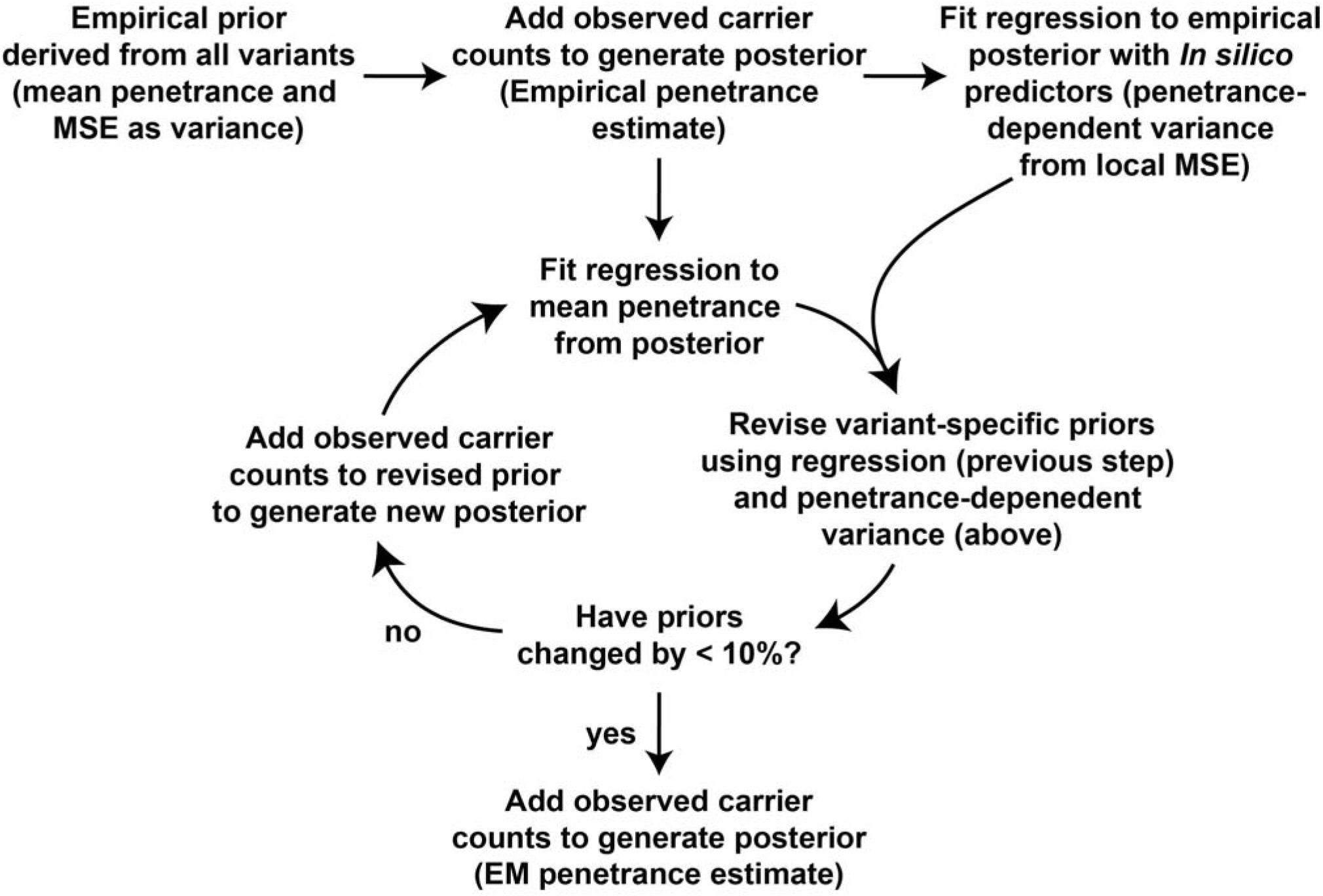
Generation of empirical and EM priors. The modified EM algorithm is an iterative technique composed of two steps: 1) calculate the expected penetrance from an empirical Bayes penetrance model and 2) fit regression of our estimated penetrance on variant-specific characteristics by maximum likelihood. The fitted model is then used to generate an updated, imputed prior and subsequent posterior expected penetrance and this process is iterated until it converges to the maximum likelihood solution, when the new priors change by less than 10% from the previous iteration.

### Collection of the SCN5A variant dataset

The dataset used was curated from 711 papers in a previous publication,^16^ to which we added an additional 45 papers on *SCN5A* that had been published since the previous dataset was constructed. Briefly, we searched publications for the number of carriers of each variant mentioned, the number of unaffected and affected individuals with diagnosed BrS, and variant-induced changes in channel function, if reported; all recorded values were normalized to wild-type values reported in the same publications. We supplemented this dataset with all *SCN5A* variants in the gnomAD database of population variation (http://gnomad.broadinstitute.org/; release 2.0).^19^ Due to the rarity of BrS (∼1 in 10,000),^20^ carriers found in gnomAD were counted as unaffected. A browsable version of the dataset, the *SCN5A* Variant Browser, is available at http://oates.mc.vanderbilt.edu/vancart/SCN5A/. We further collected *in silico* pathogenicity predictions from four commonly used servers: SIFT,^21^ Polyphen-2,^22^ CADD,^23^ and PROVEAN.^24^ We also include basic local alignment search tool position-specific scoring matrix (BLAST-PSSM) for *SCN5A*^25^ and the per residue evolutionary rate,^26^ previously shown to have predictive value for predicting functional perturbation for the cardiac potassium channel gene *KCNQ1*,^27^ and point accepted mutation score (PAM).^28^ Additionally, we leveraged structures of the *SCN5A* protein product and derived a penetrance density as previously described in an application to predict functional perturbation (see supplemental for details).^12^ We included in-frame deletions and insertions, but removed nonsense and synonymous variants from the dataset to avoid complications with the structure-based feature. This process yielded a total of 1,415 variants with at least 1 observed carrier for the analysis described below.

### Initial Empirical Bayes beta-binomial prior penetrance calculation

Using the data from the aforementioned literature curation,^16^ we estimated the penetrance for each observed variant using a beta-binomial empirical Bayes model. To estimate prior distributions for the penetrance we calculated α and β by finding the weighted mean penetrance over all variants in the dataset and estimating the variance. Weighting was done using the equation

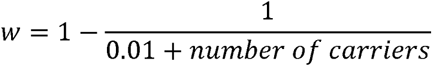

to ensure variants with a greater number of carriers, and therefore greater certainty in penetrance, had a greater weight in the preliminary analysis; number of carriers were equal to or greater than one. We estimate the variance in penetrance as the mean squared error (MSE) in this step. This derived prior distribution for penetrance, α_prior_ and β_prior_ of 0.46 and 2.71, respectively, was applied to each observed variant. The variant-specific posterior was derived by adding observed carrier counts of affected and unaffected to α_prior_ and β_prior_, respectively, and the resulting posterior mean penetrance was used in development of the subsequent predictive model (equation 1), and as one of the final estimates of posterior mean penetrance (empirical) for comparison with the EM results described below.

### Estimating variance in posterior penetrance

Modeling of estimated penetrance using previously published classifiers/predictors by linear regression was used to generate a distribution of plausible penetrance values for each variant in the dataset; penetrance predictions were truncated at 0 and 1. Penetrance dependent variance was calculated by averaging model MSE of the 100 variants with predicted penetrance closest to predicted penetrance of the variant of interest (Figure S2). Due to uncertainty in the estimated penetrance, all subsequent models and Pearson R^2^ calculations were weighted by the inverse variance of the estimated posterior beta distribution capped at the ninth decile determined in this step.

### Expectation maximization Bayesian beta-binomial penetrance predictions

The final step of this procedure employs a set of models built with a linear regression pattern mixture algorithm, updating and predicting posterior mean penetrances iteratively until the resulting α_prior,EM_ and β_prior,EM_ parameters combined changed by < 10% from the previous iteration, with a maximum of 10 iterations to circumvent oscillations. This process typically converged within three to four iterations. Some variants were estimated to have a penetrance and penetrance-dependent variance incompatible with a beta distribution; we applied the empirical prior (described above) in these cases. These final posterior estimates (EM) are used as the conditional prior for when new data become available for known variants or new variants discovered in a population.

### Assessment of predictions

To assess discrimination of predictors we calculated penetrance dependent c-statistics (AUCs) and coefficient of determination (Pearson’s R^2^); we additionally calculated frequencies of posterior mean penetrances (derived from both empirical and EM priors) within EM imputed prior 95% credible intervals. To compare the classification performance of the various predictors, we compute AUCs for two subsets of variants: all predictors are available or structure is available but peak current is not available. The two remaining subsets of variants (peak current is available but no structure or neither peak current nor structure are available) had few variants with greater than 20% posterior mean penetrance, therefore these subsets are not presented. We also report coefficients of determination weighted by the inverse variance of the estimated posterior beta distribution as described above. We used leave-one-out cross-validation (LOOCV) with the same EM iterative procedure as outlined above to further validate coefficients of determination. Furthermore, we report coefficients of determination of mean posterior penetrance (derived from either the EM prior or empirical prior) explained by the EM prior estimated mean penetrance. The data, analytic methods, and study materials have been made available to other researchers for purposes of reproducing the results or replicating the procedure. All analyses were done using the datasets provided in supplemental material and at [website to be assigned at time of publication].

## Results

Using a Bayesian beta-binomial method, we calculated a set of imputed expectation maximization priors with an interval of credible penetrance values (Figure S3). Affected and unaffected carriers (equivalent to the likelihood function) were then added to the prior to generate the posterior penetrance estimate and associated credibility interval. We applied this approach to a previously generated dataset of *SCN5A* features and BrS phenotype counts,^16^ supplemented with reports in the literature published within the last year. The penetrance estimates generated from predictive features (function, in silico predictions, structural metric) have different mean values (low for L1308F and high for R878C) as well as regions of credibility (narrow for L1308F and wide for both I1660V and R878C, Figure 2). There was a general trend that variants predicted to have relatively low penetrance had relatively narrow credible intervals compared to variants with relatively high predicted penetrance. This follows from the estimates of predicted penetrance-dependent variance which is greater as the predicted penetrance increases (Figure S2).

**Figure 2.**
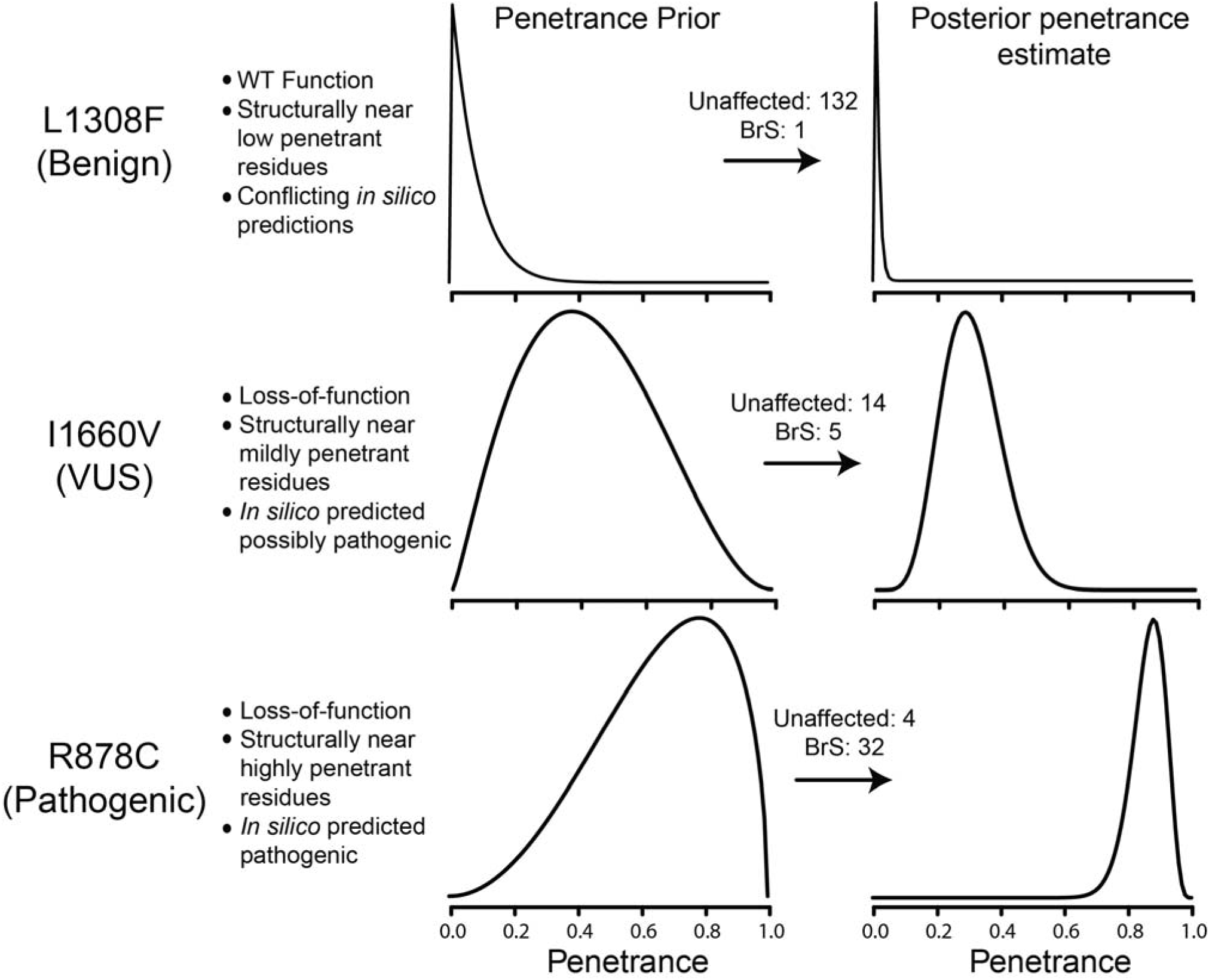
Penetrance priors are informed by variant-specific features. Probability density (y-axis) versus penetrance (x-axis) for three selected *SCN5A* variants where structure, function, and *in silico* classification are known. Numbers of affected and unaffected individuals reported are presented for each variant. The penetrance posteriors match classification as presented in ClinVar, in parentheses below variant identity. When variant-specific data are known, the penetrance estimate is adjusted to reflect the penetrance probability consistent with variants with similar features. The classification of I1660V is VUS in ClinVar;^36^ however, as an important distinction between our proposed methodology and the classification framework commonly used, we suggest *SCN5A* I1660V should be classified as a variant with close to 25% BrS penetrance.

### A modified Bayesian approach to generate priors

A typical Empirical Bayes approach would combine information across all variants to estimate a *single* prior distribution and estimate penetrance from that prior. These estimates assume all variant effects have the same prior and are therefore shrunk towards a global mean across all variants. Here we put forward a method to model the latent penetrance (mean and variance) for *each variant* from variant-specific predictive features to impute a variant-specific prior, which we then use to compute the posterior penetrance.

### Continuous prediction of penetrance: function and structure improve accuracy

We next determined which features used to predict estimated penetrance contributed to the variance explained. To accomplish this, we used EM priors to predict posterior penetrance generated from empirical priors (non-variant specific) or EM priors (variant-specific, imputed priors). The subset of all variants where function and structure are known has the highest variance explained of any subset in the pattern mixture-predicted models; the subset of variants where structure is known and peak current is unknown is similarly well predicted (Tables 1 and 2; Figure 3). The Pearson’s R^2^ is near 0.5 for both of these subsets. Leave One Out Cross Validation (LOOCV) estimated optimism is less than 0.10. These performance metrics are much improved compared to using sequence-derived predictive features alone (Pearson’s R^2^ of 0.16). Even the most conservative Pearson’s R^2^, from EM prior predictions compared against empirical estimates of mean posterior penetrance, are near 0.4 for the subset of variants where function and structure are known (and an estimated optimism of less than 0.1). All these data support the use of structure and function in estimating variant-specific penetrance, especially structural data.

**Table 1.** Weighted R^2^ from models built with the subset of variants where peak current, *in silico* predictions, and structure are known, trained and evaluated with displayed subsets of features

| Features | Empirical<sup>†</sup> | EM<sup>†</sup> |
| --- | --- | --- |
| Function | 0.13 [0.06-0.22; 0.13] | 0.16 [0.09-0.26; 0.16] |
| Function and Structure | 0.35 [0.21-0.51; 0.35] | 0.47 [0.32-0.62; 0.47] |
| Function, Structure, Seq. | 0.39 [0.26-0.52; 0.39] | 0.51 [0.38-0.63; 0.51] |
| Sequence | 0.09 [0.06-0.12; 0.09] | 0.15 [0.12-0.19; 0.15] |

<sup>†</sup>Adj. $R^2$ [95% Confidence Interval; LOOCV]

**Table 2.** Weighted R^2^ from pattern mixture model evaluating subsets of all variants in the database

| Variant Subset | Empirical<sup>†</sup> | EM<sup>†</sup> |
| --- | --- | --- |
| Whole dataset | 0.22 [0.15-0.31; 0.20] | 0.44 [0.36-0.51; 0.41] |
| Function, no Structure | 0.03 [0.00-0.14; 0.03] | 0.05 [0.00-0.20; 0.05] |
| Function and Structure | 0.41 [0.27-0.55; 0.33] | 0.54 [0.41-0.67; 0.45] |
| No Function or Structure | 0.00 [0.00-0.02; 0.00] | 0.16 [0.09-0.25; 0.15] |
| Structure, no Function | 0.20 [0.10-0.35; 0.19] | 0.47 [0.35-0.59; 0.46] |

<sup>†</sup>Adj. $R^2$ [95% Confidence Interval; LOOCV]

**Figure 3.**
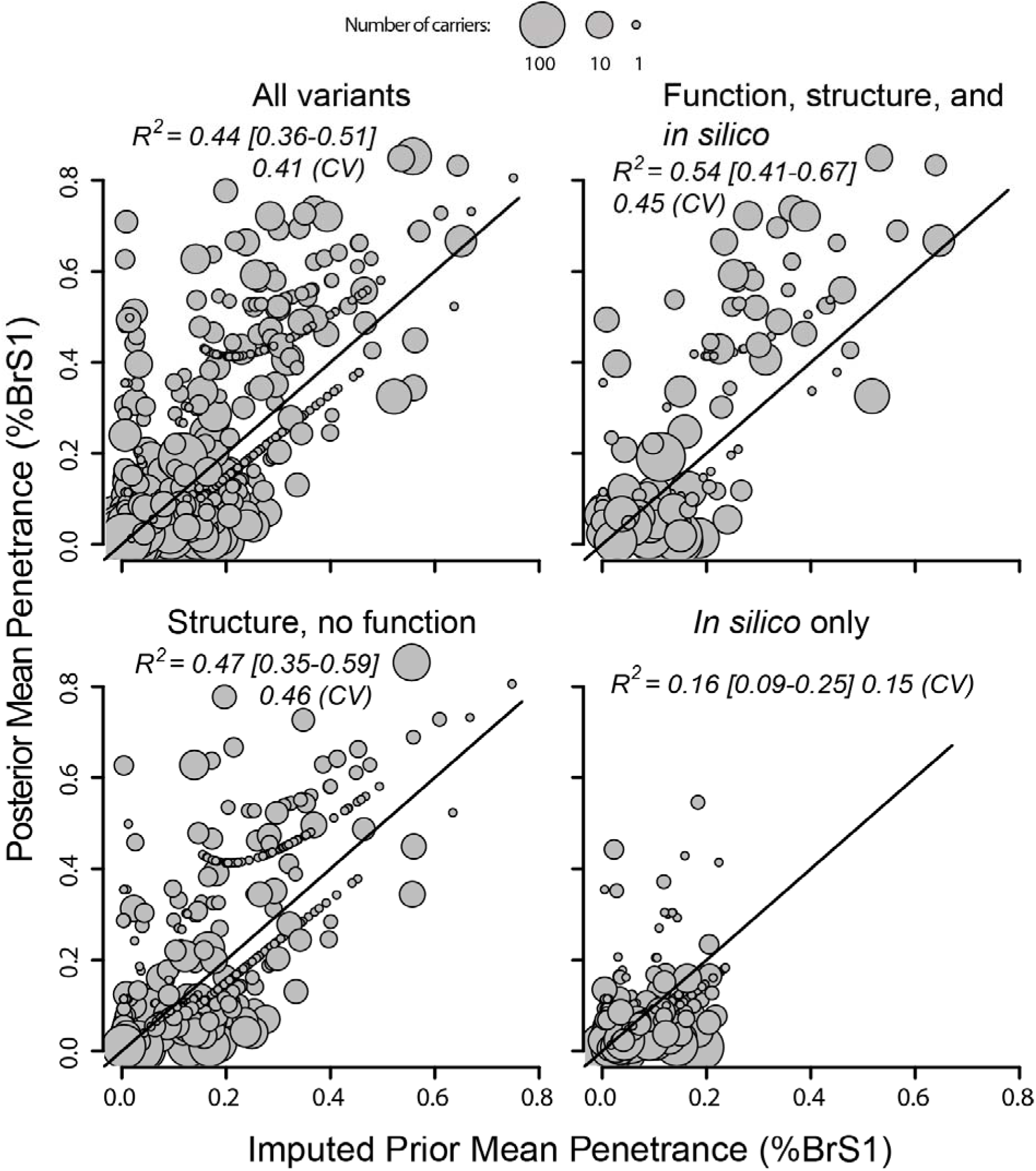
Significant variation in mean posterior BrS penetrance is explained by function and structure of *SCN5A* variants. Mean penetrances from the EM pattern mixture model for all variants before (imputed) and after (posterior) affected/unaffected carrier count was added. Variants are subsetted according to the text above the figure.

### Classification of high or low penetrance variants

One important application of classifying variants is to focus resources on variants most likely to cause some pathology or potentially explain an existing pathology. To this end, we employed a binary classification (disease-causing vs. not disease-causing), selecting multiple penetrance cutoffs. Since variants varied by which predictors were available, we divided variants into subgroups and attempted classification within those groups. The subgroups were 1) function and structure are known and 2) structure is known, but function is unknown. The sharp rise on the left side of the ROC curves in Figure 4, suggest function (peak current) and structure are best at discriminating low from high BrS penetrance variants, and that this classification performance is best for variants predicted to be most highly penetrant (i.e. those with more severe in vitro phenotype and close in space to other variants with high BrS penetrance). The gap between ROC curves for models trained with all features (blue) and each feature individually suggests the non-overlap of information contained in any one feature, most notable in the upper right figure (structure available but no peak current).

**Figure 4.**
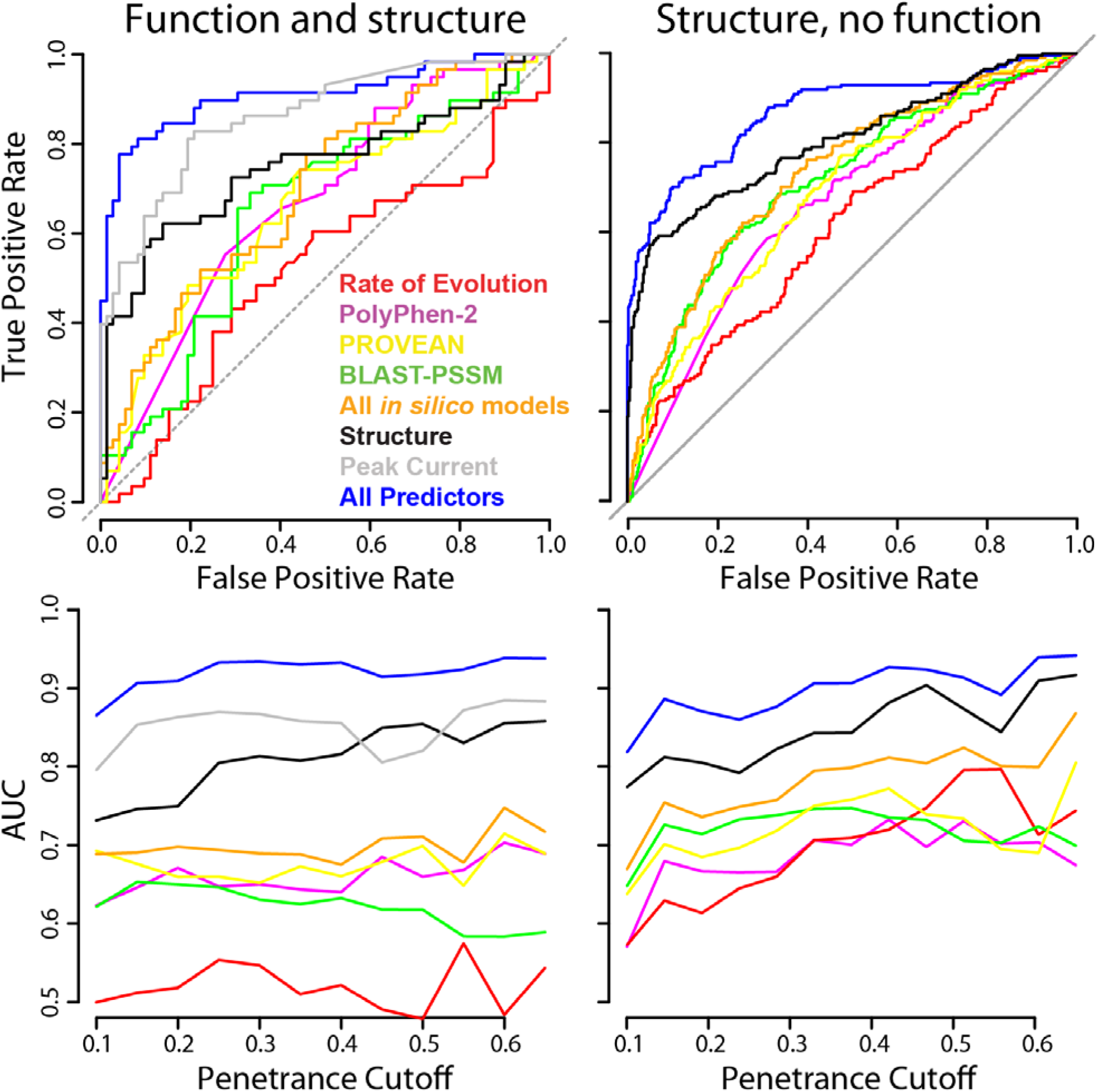
Function and structure inform variant classification. Above: ROC curves classifying high versus low penetrance; below: AUCs determined at multiple penetrance cutoffs. Number of variants in each subgroup are the following (number pathogenic at a cutoff of 20%): 130 (58) and 623 (201) for variants with peak current and structure or variants with structure and no peak current, respectively. Here *in silico* models refer to previously published predictive classifiers, as described in the methods.

### Penetrance prediction precision

Our goal is to generate more accurate predictions of variant-specific penetrance and also to quantify the uncertainty of our predictions to yield an informative and interpretable priors and posteriors. One concern with the proposed methodology, indeed much of Bayesian prediction, is that posterior mean estimates of penetrance are derived from observations of affected and unaffected individuals (likelihood function), but are also influenced by priors. We believe the most accurate prior is imputed by variant specific maximized likelihood (EM prior); however, for variants with low carrier counts, the resulting posterior mean penetrance estimates are determined largely by the prior. To measure the optimism in our estimates of precision, we calculated frequency statistics of posterior mean penetrance estimates derived from the empirical prior or imputed EM prior falling within 95% credible intervals from pattern mixture EM priors. If variant-specific EM priors were overly optimistic in precision, we expect the fraction of posterior mean penetrances within 95% credible intervals to fall below 95% as we select variants with greater carrier counts (i.e. less influenced by the EM prior) for evaluation; the effect would be especially noticeable when using the empirical prior to generate posterior mean penetrances. However, we observe the frequency at which posterior mean penetrance estimates fall within the 95% credible interval defined by the prior is at or exceeds 95% (Figure S4). As expected, the percentage is the lowest for low carrier count variants with a posterior mean penetrance derived from an empirical prior, though still nominally at 95%.

## Discussion

### Estimates of penetrance are informed by variant-specific features

A key assumption of the methodology proposed here is that variants with similar determinable features, such as function and evolutionary sequence conservation, have similar penetrance characteristics—we suggest properties extrinsic to *SCN5A* variant identity are responsible, at least in part, for the BrS penetrance observed clinically. As an example, nonsense variants or variants that produce no sodium current, result in non-negligible penetrance of BrS.^16^ At the other extreme, variants that have WT characteristics and low sequence conservation lead to negligible BrS penetrance. We propose here a Bayesian beta-binomial framework to contain both of these extremes as boundary conditions and a continuous estimate between the two. We put forward this methodology as a means to compare variants with greater total carrier counts to rare variants with less available data along the continuous axis of penetrance. The resulting penetrance and uncertainty estimates yield a variant-specific prior interpretable by clinicians and carriers of these variants as equivalent to hypothetical observations of affected and unaffected carriers (α_prior_ and β_prior_, respectively).

### Estimates of penetrance are more informative than variant classification

Here we develop a method to estimate penetrance by quantitatively incorporating variant-specific information into a probabilistic framework. While clinical evidence affirms a strong relationship between *SCN5A* variants and BrS, many genetic and environmental factors influence the ultimate presentation of BrS in an individual (Figure 5).^29-31^ Some variants affect almost all known carriers and some variants confer only modest increased risk.^14,15^ One lesson from our previous analysis of *SCN5A* function and its influence on penetrance was that, without exception, any well-sampled variant (more than 30 carriers) had at least one individual without any known arrhythmia phenotype—no variant is 100% penetrant.^16^ This suggests the categorical ACMG terms (e.g. “pathogenic”, “benign”) are not the optimal way to describe the impact of genetic variants on disease. We instead suggest carrier risk is probabilistic and lies on a continuous range from 0 (impossible) to 1 (certain), mirrored in the estimation of penetrance.

**Figure 5.**
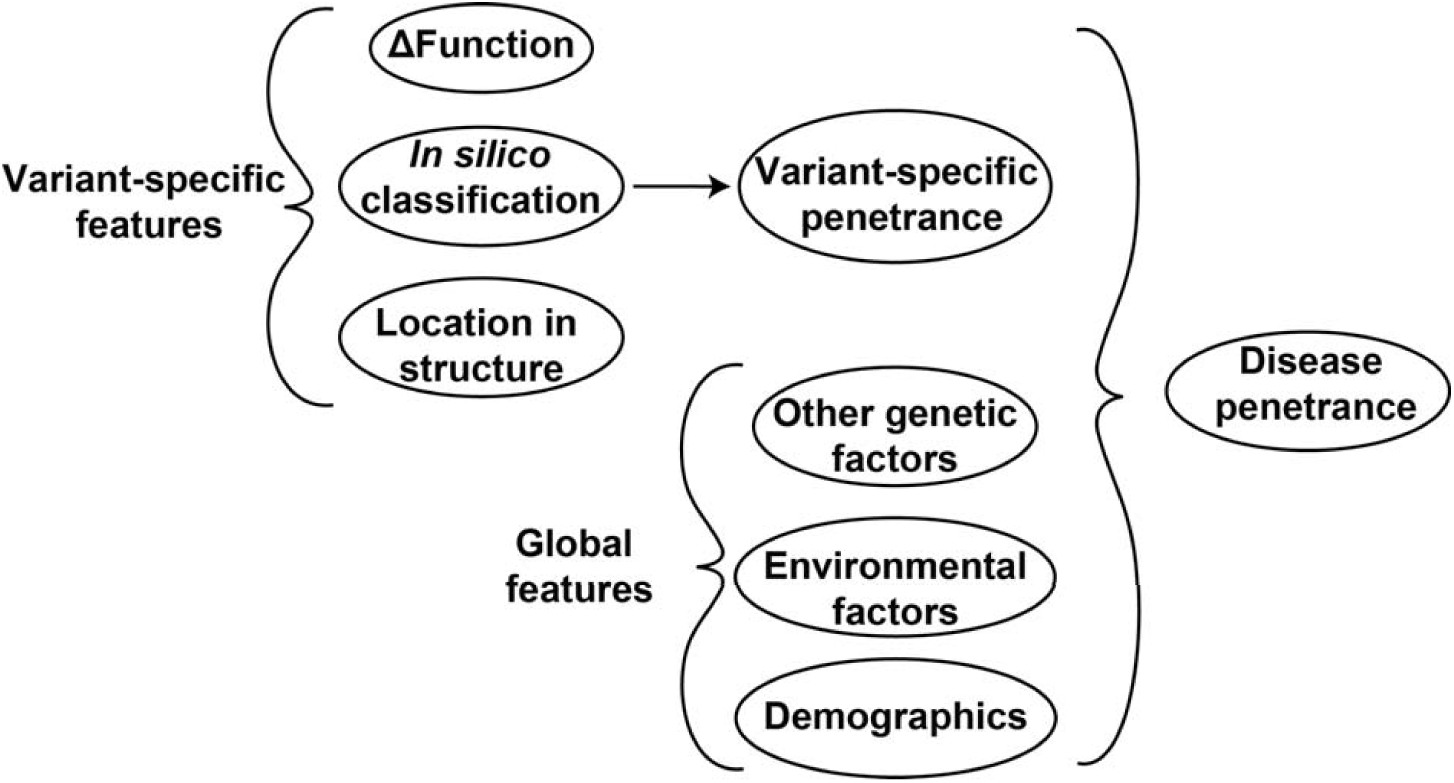
Factors influencing disease presentation. The objective of this paper is to quantitatively address the relationship between variant-specific features and the variation in penetrance attributable to specific variants. Given the observed relationship between penetrance and variant-specific features, we suggest BrS penetrance decomposed into variant-specific features and other contributing factors.

### Structure and peak current improve prediction of penetrance

Here we showed that variants with increased penetrance burden of BrS tend to localize in protein structure (Figure 6 and Tables 1 and 2). Features derived from structure contain information not present in other predictive features, as can be seen by the improvement in prediction when structure is included, true for all subsets evaluated (Tables 1 and 2 and Figure 3). The degree of information added by structure suggests the three-dimensional location in regions enriched for higher penetrance do not also have functional disruptions or evolutionary constraints, as encoded in peak current and sequence-based predictive features, respectively. One potential explanation is that the functional perturbation used, peak current, imperfectly recapitulates the functional defect responsible for variation in penetrance, or perhaps only a subset of mechanisms that result in lower peak current have a large influence on BrS clinical presentation (akin to what we have observed with late current and long QT syndrome^32^). Another possible explanation is peak current contains noise from the variability in measurements from different labs or different model cell systems which dilutes the otherwise observable relationship between loss of peak current and BrS penetrance. Whatever the reason, clearly there is a need to include structural information in variant interpretation.^33-35^

**Figure 6.**
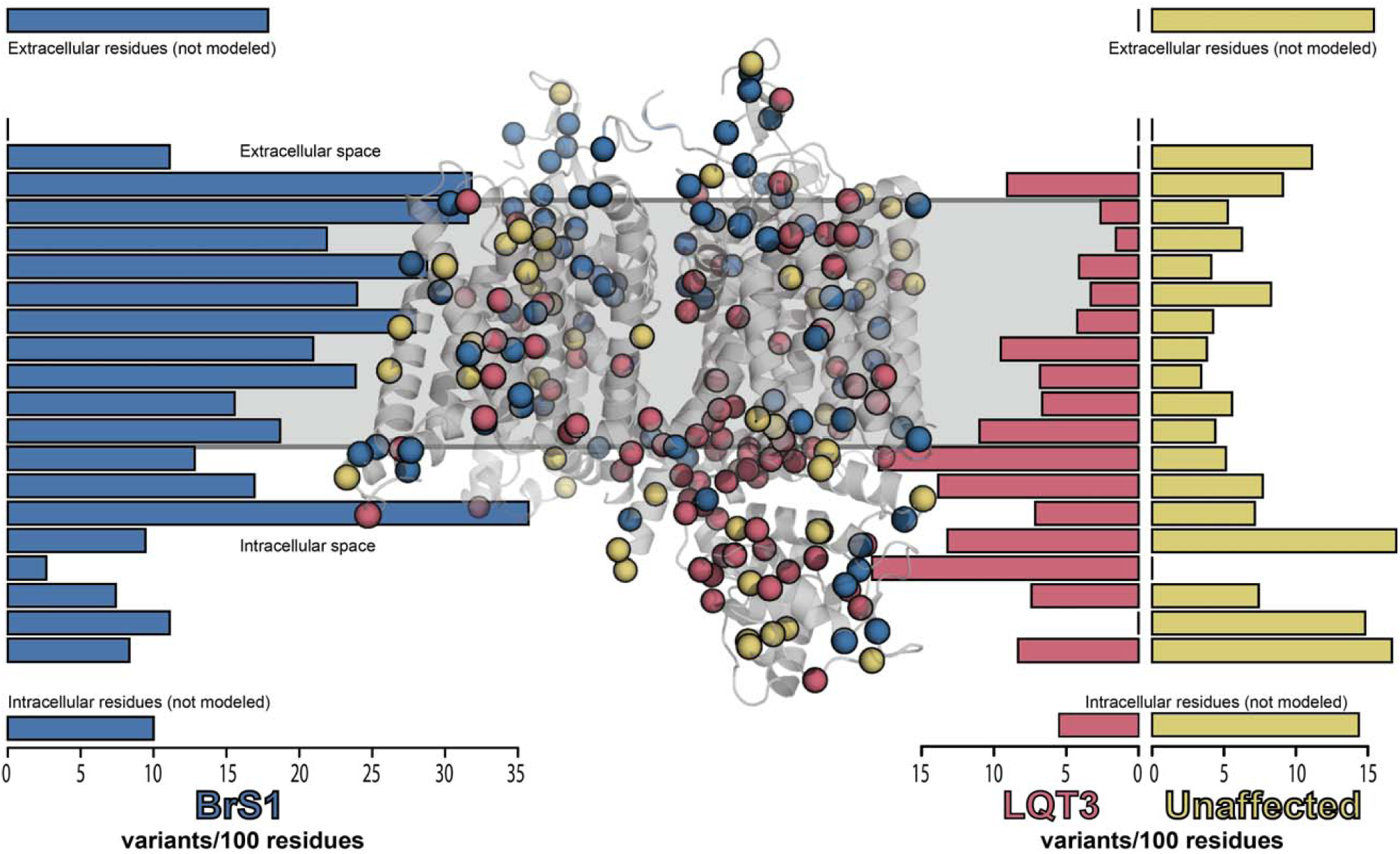
*SCN5A* pathogenic and benign variants cluster in space. Rate of variants associated with BrS (blue) or LQT3 (red) or from unaffected populations (control, gold) in a model of the *SCN5A* protein product transmembrane domain. Each bar represents a histogram bundling of variants within a 5Å window in three-dimensional space, boxes at each of the four corners represent residues not modeled (only 31 residues were not modeled in the extracellular loops). There is a relative paucity of control variants within the structured transmembrane region and the relative abundance of both BrS and LQT3 in the same region. LQT3 variants are more frequent in the intracellular half of the model, and BrS variants are more frequent in the extracellular half. The membrane-flanking, intracellular part of SCN5A controls the inactivation which, when compromised, frequently results in LQT3. The BrS enrichment near the extracellular half is likely due to more compacting of residues in the top half of the pore domain, more often leading to a destabilizing influence of amino-acid substitution.

### Prospects for applications of this method

The methodology described relies upon having a sufficient number of variants with high carrier counts such that penetrance can be reliably estimated and also having predictive features with some relationship to the disease (e.g. changes in function and sequence conservation). This limits the potential application of the methods described herein to a relatively small subset of genes at present. For example, of the 59 genes the ACMG recommends clinical diagnostic laboratories report secondary variant discovery, 36 have greater than or equal to 20 missense “pathogenic”/”likely pathogenic” variants in ClinVar,^36^ suggesting that many variants are described in the literature and can be curated in a similar manner to *SCN5A*.^36^ The penetrance estimates in our approach will continue to be refined as additional data become available (i.e. phenotype data from case reports and large biobank projects, additional *in vitro* functional studies, and improved computational and structural predictors).^13,30,37-39^

## Conclusion

Penetrance, as formulated in a Bayesian beta-binomial framework, allows us to quantitatively integrate phenotypic data with functional measurements, variant classifiers, and sequence-and structure-based features to accurately estimate disease risk attributable to specific variants, even when clinical information is limited. Penetrance more precisely describes disease risk and uncertainty than categorical pathogenicity classifications. We suggest this probabilistic penetrance approach can be applied to additional rare Mendelian diseases to better estimate disease risk and improve the impact and accuracy of genomic medicine.

Collected data are available at the following website: (to be finalized with publication)

## Acknowledgments

Christian M. Shaffer for analysis and help building the website. This research was funded by R00HL135442 (BMK), P50GM115305 (DMR), and F32HL137385 (AMG).

