## Supplement for "A Bayesian method using sparse data to estimate penetrance of disease-associated genetic variants"

### **Supplemental Methods**

We calculated BrS penetrance densities based on an approach akin to k-nearest neighbors. We calculated penetrance density by averaging empirical BrS mean posterior penetrance of variants near the variant of interest weighted by the inverse of their distance from the variant of interest. This calculated feature therefore depends on how many variants are near the variant of interest, with regions in three-dimensional space dense with high penetrance variants—“hotspots”—yielding a higher penetrance prediction. Penetrance density is calculated as follows:

$$\rho_{j}=\frac{\sum_{i=0}^{n} {BrS Posterior Penetrance}_{i,mutation\left( i\neq j \right)}\cdot\frac{1}{1+e^{(\frac{d_{i,j}}{2})}}}{\sum_{i=0}^{n} \frac{1}{1+e^{(\frac{d_{i,j}}{2})}}}$$

where ρ_j_ is BrS penetrance density of the j^th^ residue, *BrS Posterior Penetrance_x_*_,i,mutation(i≠j)_ is the penetrance of BrS for the i^th^ variant, and d_i,j_ is the distance between the center of mass of residues i and j. i does include residue j, but only if the identity of the amino-acid mutation is changed, i.e. mutation(i) ≠ mutation(j).

We previously evaluated many structure-based predictive features and found only one, “functional density”, here “penetrance density”, calculated as a three-dimensional cousin to k-nearest neighbor, to have consistent additive value and so we include that feature termed “structure” for simplicity throughout the text.

*Influence of variants with small carrier counts.* It is difficult to know *a priori* how to weight variants with only a single observation in either an affected or unaffected population; and this is the class of variant with the most members. However, differences in performance mentioned suggest non-negligible contribution to Pearson’s R^2^ by variants with only a single known carrier (Tables 1 and 2). If we look only at variants with five or more known carriers, the Pearson’s R^2^ from the LOO cross validated empirical posterior predictions increase from 0.20 to 0.28 for the subset of variants where structure is known but function is unknown. Given the influence of the prior in BrS posterior penetrance estimates for variants with few carriers, these differences suggest the inclusion of single-carrier variants make models evaluated by empirical mean posterior estimates more pessimistic but also possibly inflate the performance metrics for models evaluated by EM estimated posteriors.

*Expected generalizability of penetrance predictions.* We chose a parsimonious regression methodology as well as relatively few features to avoid overfitting. The fraction of variants whose posterior mean (derived from either the empirical or EM prior) falls within the EM prior penetrance predicted 95% credible interval is 95% or better, even when stratified by carrier count. Thus, the resulting data-driven EM beta-binomial prior distributions are not overconfident. Indeed, the EM generated prior penetrance distributions may be too conservative in variance. However, since variants with known structure and function come from a biased subset of all variants, we speculate the coefficient of determination values will generalize somewhere between those observed for all variants using the pattern mixture model and the subset of variants where structure and peak current are known.

### **Figure Legends**

Figure S1. BrS penetrance probability versus penetrance for the empirical prior.

Figure S2. Penetrance variance estimate. Posterior penentrance prediction mean squared error (y-axis) versus penetrance prediction (x-axis). The solid line indicates a nearest 100 variants average, which is what we used to estimate variance in penetrance estimates (see methods).

Figure S3. Histogram of BrS penetrance imputed EM prior posterior means and associated upper and lower bounds to 95% credible interval from pattern mixture models. Plotted are BrS posterior mean penetrances from imputed EM priors (“Predicted”, green) and upper (red) and lower (blue) bounds to associated 95% credible intervals from those imputed EM priors.

Figure S4. Fraction of variants with posterior mean penetrance values that fall within imputed EM penetrance prior 95% credible intervals. Two sets of posterior penetrance values were used, one derived from empirical prior and carrier observations and another derived from imputed EM prior and carrier count observations. A carrier count cutoff was used to select variants with at least the number of carriers indicated on the x-axis for this frequency analysis.

Figure S5. Histograms of EM pattern mixture penetrance prior α (or hypothetical affected carrier count) and β (or hypothetical unaffected carrier count) for all variants.

### **Figures**

Figure S1

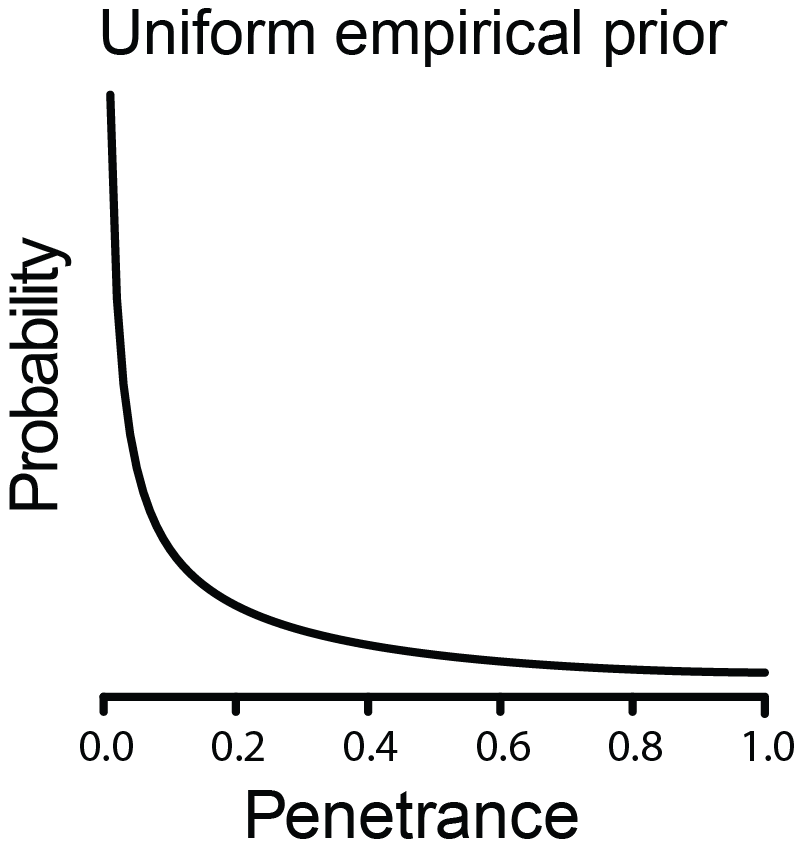

Figure S2

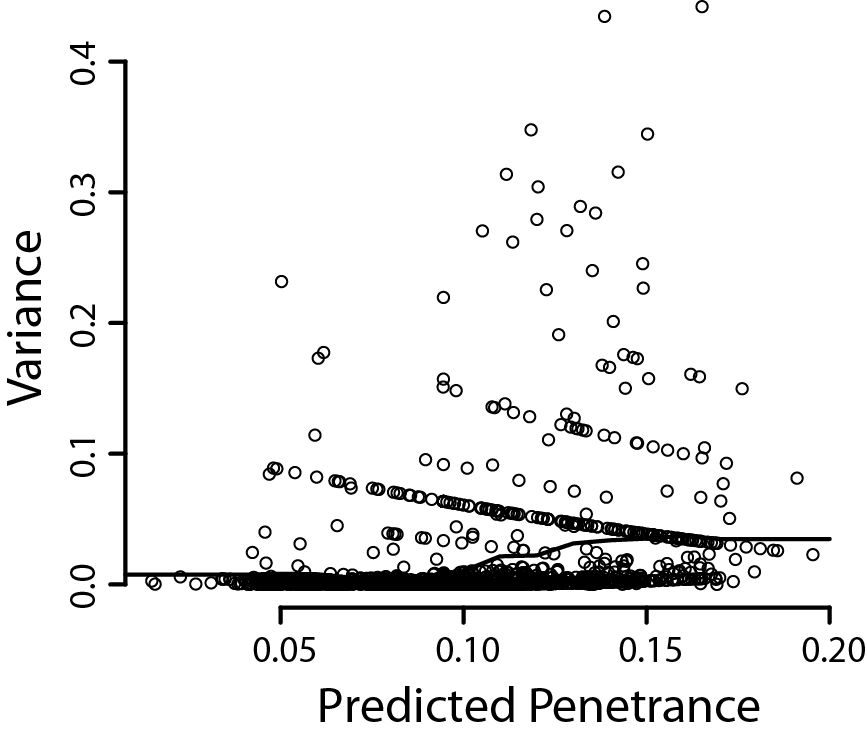

Figure S3

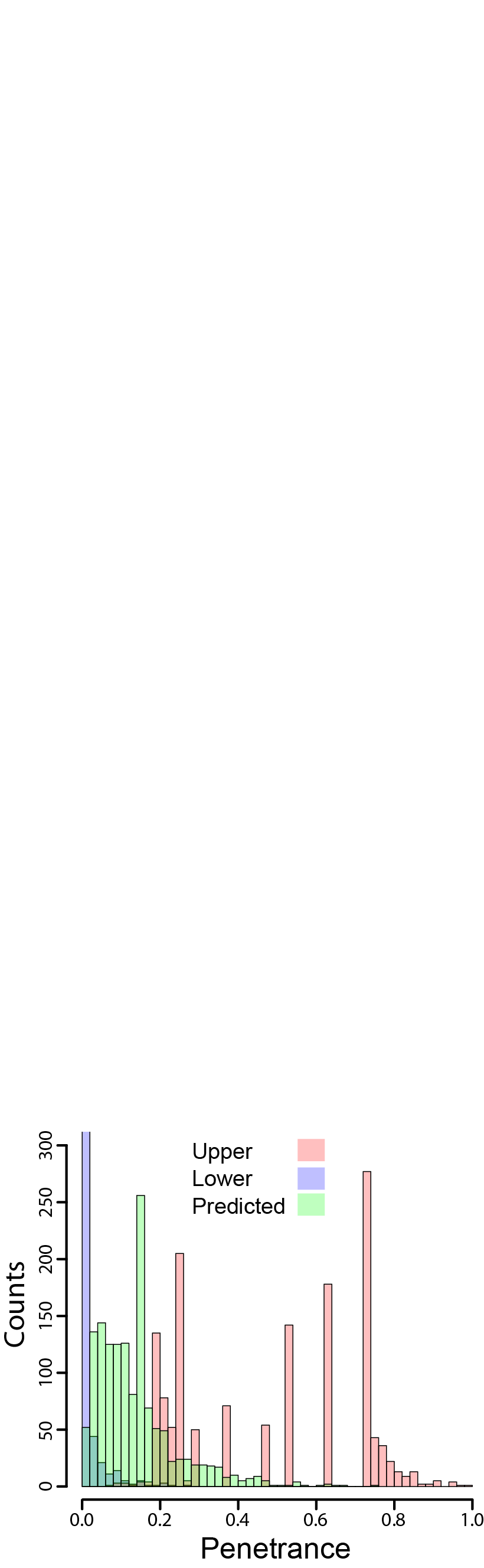

Figure S4

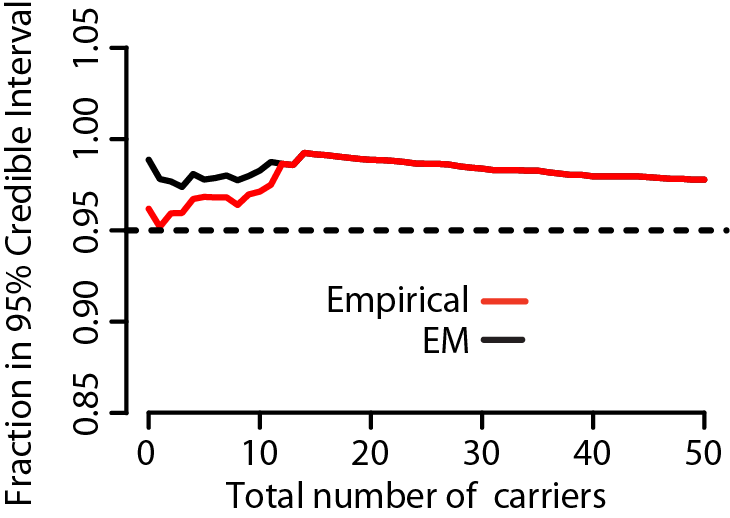

Figure S5

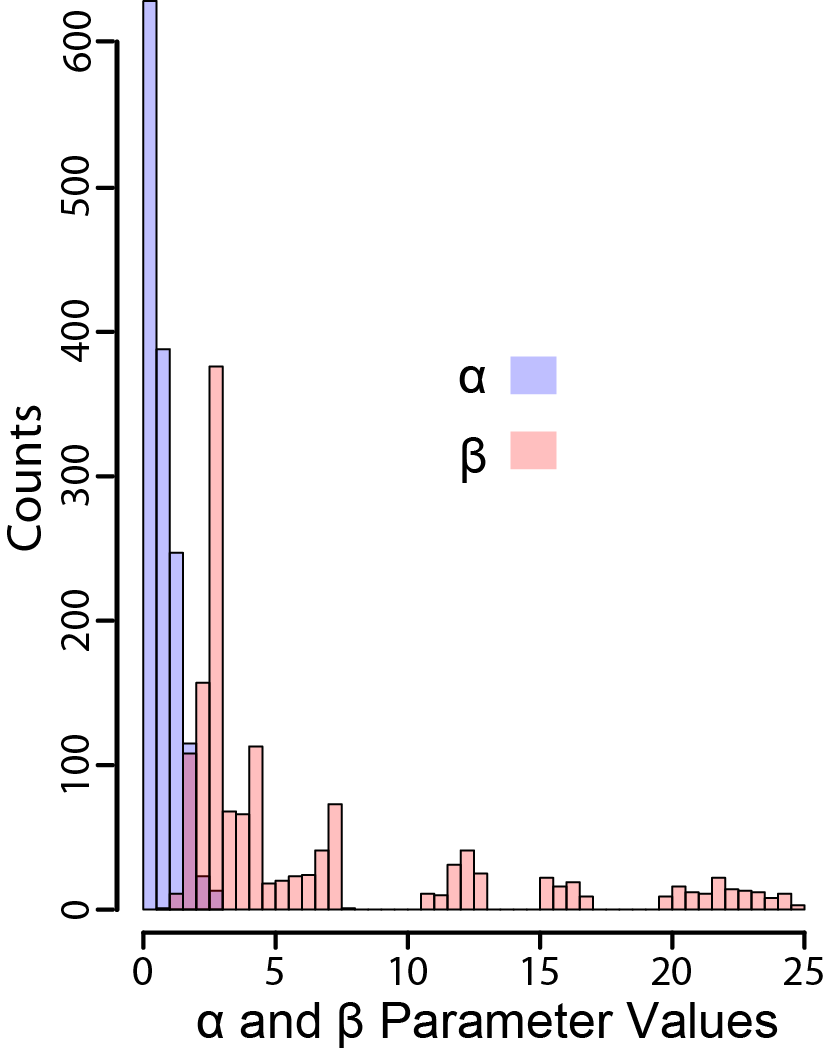
